# Regulation of Decay Accelerating Factor primes human germinal center B cells for phagocytosis

**DOI:** 10.1101/2020.08.27.271148

**Authors:** Andy Dernstedt, Jana Leidig, Anna Holm, Jenny Mjösberg, Clas Ahlm, Johan Henriksson, Magnus Hultdin, Mattias NE Forsell

## Abstract

Germinal centers (GC) are sites for extensive B cell proliferation and homeostasis is maintained by programmed cell death. The complement regulatory protein Decay Accelerating Factor (DAF) blocks complement deposition host cells and therefore also phagocytosis of cells. Here, we show that B cells downregulate DAF upon BCR engagement and that T cell-dependent stimuli preferentially led to activation of DAF^lo^ B cells. Consistent with this, a majority of light and dark zone GC B cells were DAF^lo^ and susceptible to complement-dependent phagocytosis, as compared with DAF^hi^ GC B cells. We could also show that the DAF^hi^ GC B cell subset had increased expression of the plasma cell marker Blimp-1. DAF expression was also modulated during B cell hematopoiesis in the human bone marrow. Collectively, our results reveal a novel role of DAF to pre-prime activated human B cells for phagocytosis prior to apoptosis.

## Introduction

High affinity memory B cells and plasma cells (PCs) have undergone two distinct selection processes; during B cell development in the bone marrow^1^ and during T cell-dependent germinal center (GC) responses in secondary lymphoid organs^2–7^. The GC is traditionally divided into two zones based on their histological appearance where proliferation and somatic hypermutation occur in the dark zone (DZ) and T cell dependent selection in the light zone (LZ)^8^.

Both the bone marrow and GCs are sites of extensive B cell proliferation. Without BCR engagement and co-stimulatory signals from CD4^+^ T cells, GC B cells undergo apoptosis and die. The clearance of apoptotic cells by phagocytosis is a critical process to maintain homeostasis and to restrict development of autoimmunity or inflammation^9,10^. Approximately 50% of all B cells die every 6 hours during a GC reaction^11,12^, and these are removed by tingible body macrophages or marginal reticular cells^13–16^. It has been suggested that human GC B cells are primed for apoptosis and phagocytosis^17^, but a mechanism for this priming has not been demonstrated.

Covalent attachment of complement components to cell surfaces is an important cue for phagocytosis^18^. This is facilitated by C3 convertase that facilitates attachement of C3b the cell surface. This process is regulated by a number of complement regulatory proteins that inhibit complement mediated phagocytosis or lysis of healthy cells^18^. The Decay Accelerating Factor (DAF or CD55) is a glycosylphosphatidylinositol (GPI) anchored protein that inhibits C3 convertase formation, and is highly expressed on B cells^19^. Nonsense mutations in the *CD55* gene leads to increased deposition of C3d on T cells and severe disease^20^. Another example of DAF-deficiency is Paroxysmal nocturnal hemoglobinuria (PNH) where some hematopoietic stem cells have defect anchoring of DAF to cell surfaces due to a somatic mutation that inhibits generation of the GPI anchor^21^. As a consequence, downstream hematopoietic cells lack GPI anchored proteins, including DAF. It was found that DAF-deficient B cells are unswitched and mainly naïve in PNH patients, whereas their DAF-expressing counterparts appear normal^22^. Since PNH patients lack all GPI anchored proteins on their DAF-deficient B cells, more targeted investigations of DAF expression on healthy B cells are required to understand if the complement regulatory protein may play a direct role in human T cell-dependent B cell responses.

Due to its critical role for inhibition of C3 convertase, we hypothesized that GC B cells regulate DAF expression to become pre-primed for phagocytosis. To test this hypothesis, we set out to investigate if regulation of DAF occurs on specific subsets of human B cells in circulation, tonsils, and in bone marrow.

## Results

### Circulating DAF^lo^ B cells are expanded during viral infection

While DAF is highly expressed on circulating B cells during steady-state^19^, we wanted to understand if DAF can be regulated by extrinsic factors such as infection. Therefore, we compared surface expression of DAF on B cells from healthy donors and from patients diagnosed with hantavirus infection and Hemorrhagic Fever with Renal Syndrome (HFRS)^23^. There, we found that naïve B cells (CD27^−^ IgD^+^), unswitched memory B cells (CD27^+^ IgD^+^), switched memory B cells (CD27^+^ IgD^−^) and CD27^−^ IgD^−^ B cells all had high surface expression of DAF in healthy individuals (Figure 1A, B). In comparison, expression of DAF was overall lower in HFRS patients, and most noticeable in the unswitched memory and the CD27^−^ IgD^−^ B cells compartments (Figure 1C). DAF expression on PBMCs did not change following short-term or long-term exposure to the causative agent of HFRS, the Puumala hantavirus (Supplemental Figure S1). Additional analyses of the DAF^lo^ CD27^−^ IgD^−^ B cells revealed that they comprised a major population of atypical B cells, characterized by low expression of the complement receptor CD21, and high expression of the inhibitory Ig receptor-like Fcrl5 and FAS-receptor CD95 during HFRS (Figure 1D). We did not observe a comparably strong phenotype on CD27^−^ IgD^−^ B cells from healthy individuals. This suggested that the infection had indirectly caused the accumulation of DAF^lo^ B cells in circulation.

**Figure 1.**
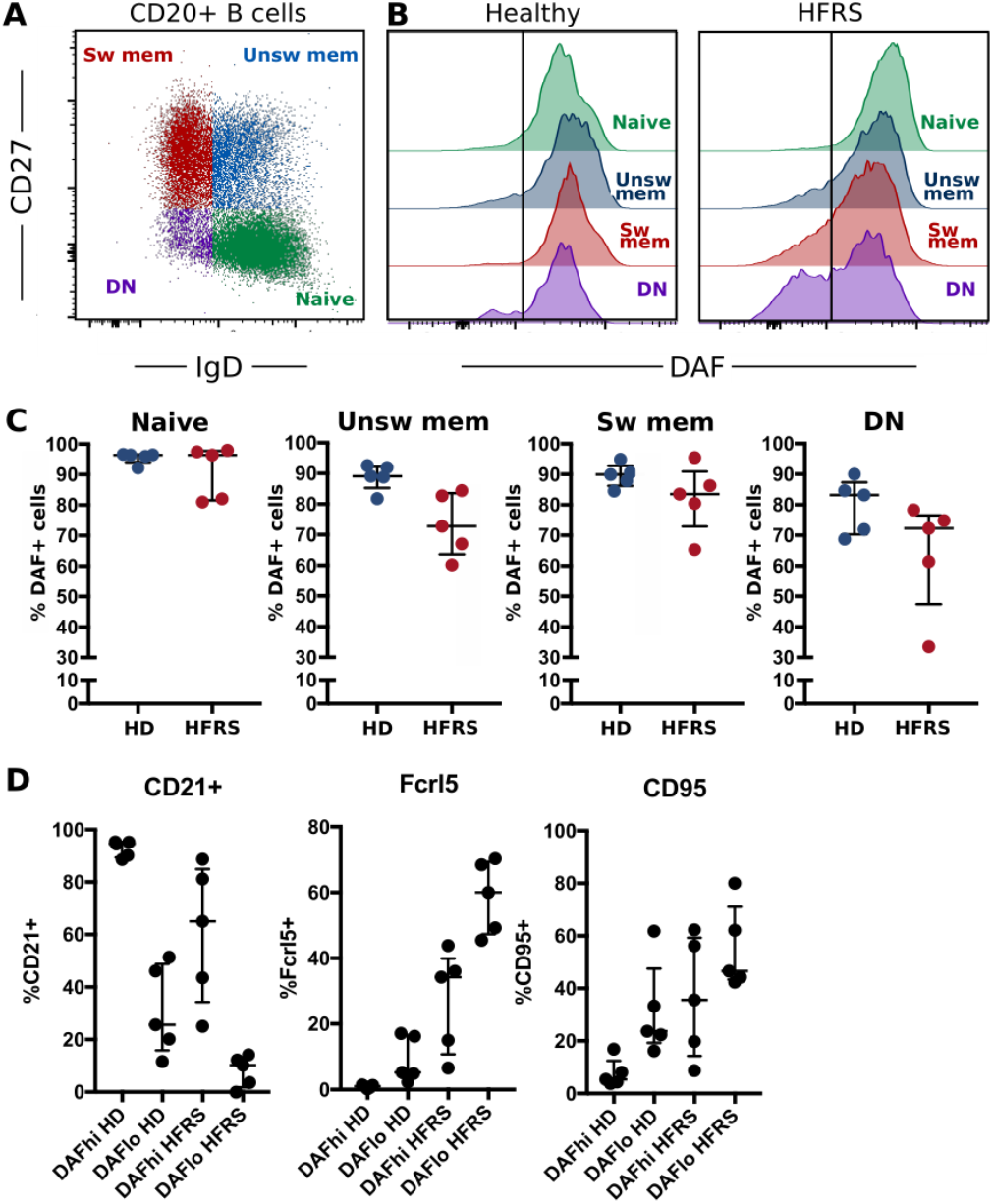
DAF expression on circulating B cells is decreased during virus infection. **(A)** Flow cytometric gating to discriminate Naïve (CD27^−^ IgD^+^), unswitched memory (unsw mem, CD27^+^ IgD^+^), switched memory (sw mem, CD27^+^ IgD^−^) and double negative (DN, CD27^−^ IgD^−^) CD20^+^ B cells. **(B)** Representative histogram plots of DAF expression on B cell subsets in circulation of healthy controls (middle) or HFRS patients (right). **(C)** Quantification of DAF^hi^ B cells in healthy donors or HFRS patients. **(D)** Expression of CD21, Fcrl5 and CD95 in DAF^hi^ and DAF^lo^ DN B cells of HD or HFRS patients. N = 5 (healthy donors) and 5 (HFRS patients).

### Specific downregulation of DAF on human GC B cells

To assess if downregulation of DAF occurs in lymphoid tissues, we performed a direct comparison of DAF expression between B cell subsets in circulation and in tonsils. There, we found that the frequency of DAF^hi^ B cells was reduced on all subsets in tonsils as compared to blood but that this reduction was most prominent in the two tonsillar IgD^−^ subsets (Figure 2A). By staining the cryosections of the corresponding tonsils, we could verify that DAF expression was downregulated in the B cell follicles, and that this coincided with low or non-detectable IgD staining (Figure 2B). In addition, low DAF detection occurred in areas of follicles that exhibited high expression of the chemokine receptor CXCR4. This suggested that DAF was specifically downregulated in GC B cells. We therefore assessed DAF expression on CD19^+^ CD20^+^ naive B cells (CD38^−^ IgD^+^), unswitched activated B cells (CD38^+^ IgD^+^), GC B cells (CD38^+^ IgD^−^), memory B cells (CD38^−^ IgD^−^), and CD19^+^CD20^+^ plasmablasts (PB) (CD38^++^ IgD^−^) from tonsils (Figure 2C). Consistent with the histology, we found that a majority of GC B cells had downregulated DAF, whereas only a minor downregulation of DAF was detected on other non-naïve subsets (Figure 2D).

**Figure 2.**
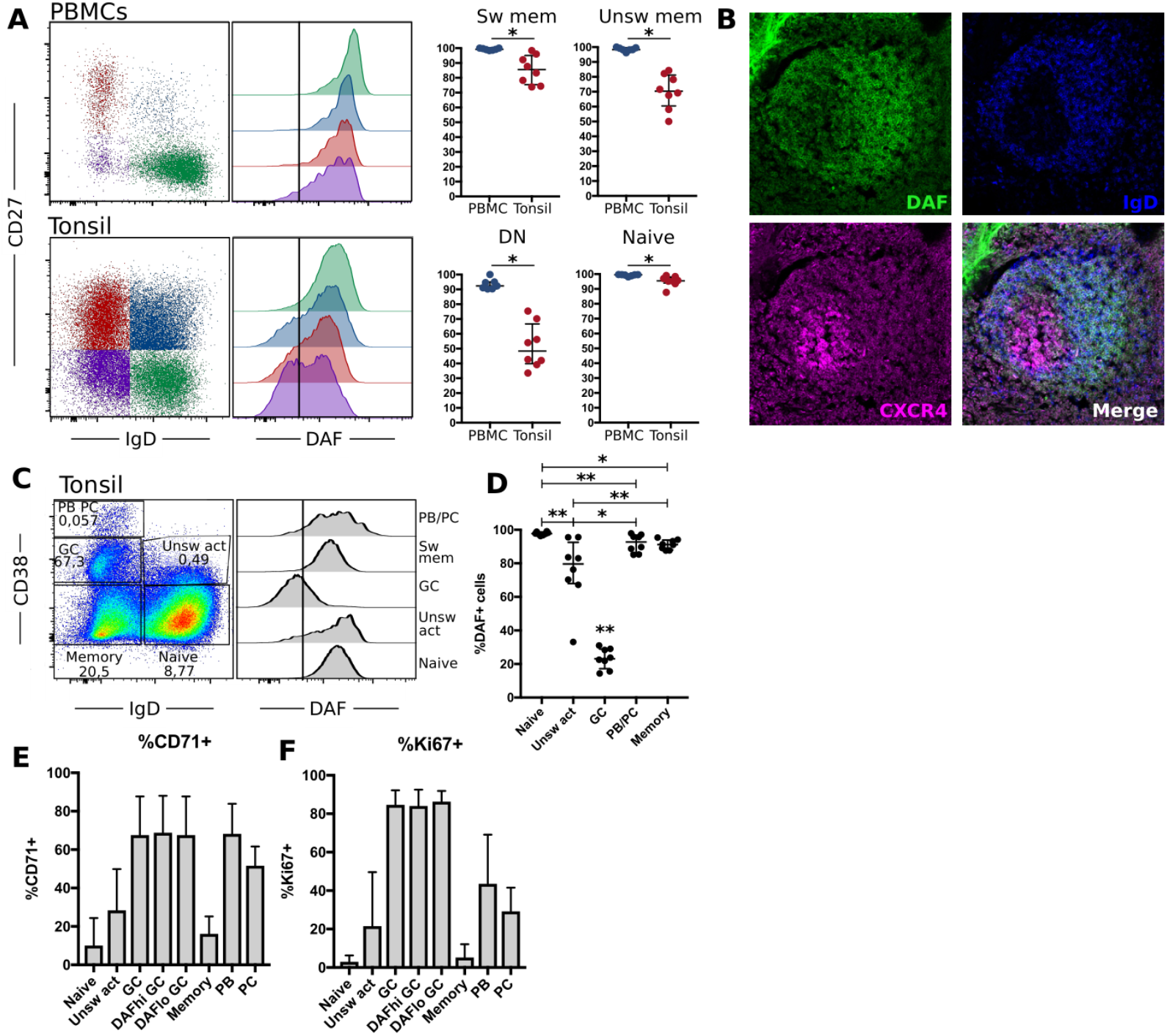
Germinal center B cells specifically downregulate DAF. **(A)** Representative flow cytometric plots and quantification of DAF expression on CD19^+^ CD20^+^ naïve (CD27^−^ IgD^+^), unswitched memory (unsw mem, CD27^+^ IgD^+^), switched memory (sw mem, CD27^+^ IgD^−^) or DN (CD27^−^ IgD^−^) B cells from circulation (N=7) and tonsils (N=8). Individual dots for each sample are shown and lines indicate median with interquartile range. **(B)** Representative tonsil section stained with DAF (green), IgD (blue) and CXCR4 (magenta). **(C)** Representative plot for subset analysis of DAF expression on germinal center cells (GC, CD38^+^, IgD^+^), naïve B cells (CD38^−^ IgD^+^), unswitched activated (unsw act, CD38^+^ IgD^+^), memory cells (CD38^−^ IgD^−^), and plasmablasts (PB) (CD38^++^ IgD^−^) of CD19^+^ CD20^+^ tonsillar B cells. Vertical line indicates the cutoff to discriminate DAF^hi^ from DAF^lo^ cells. **(D)** Quantification of DAF expression on tonsillar B cell subsets. Individual dots for each sample are shown and lines indicate median with interquartile range. **(E)** Frequency of B cell subsets with high expression of the transferrin receptor CD71. Data is represented as bar graphs showing median and interquartile range. **(F)** Frequency of B cell subsets with high expression of Ki67. Data is represented as bar graphs showing median and interquartile range. *P < 0,05; **P < 0,01 by Wilcoxon signed-rank test.

By measurement of the cell cycle activity protein Ki67 and the transferrin receptor CD71, we found that both DAF^hi^ and DAF^lo^ GC B cells demonstrated a comparably high level of activation in contrast to naïve, unswitched or swithched memory B cells (Figure 2E, F). We could also identify that DAF^hi^ GC B cells had higher expression of both CD21 (Figure 2G) and CD95 (Figure 2H) as compared with DAF^lo^ GC B cells. Since CD21 has been shown to reduce the threshold for BCR stimulation, and CD95 is critical for the extrinsic apoptosis pathway, this suggests that DAF^lo^ GC B cells have reduced potential for BCR stimulation and are less sensitive to apoptosis than their DAF^hi^ counterparts^24,25^.

### Transcriptomic profile of DAF^hi^ and DAF^lo^ GC B cells

Next, we assessed the transcriptional profile of bulk sorted DAF^hi^ and DAF^lo^ GC B cells by a Human Clariom-D microarray. Analysis of resulting data revealed that both DAF^hi^ and DAF^lo^ GC B cells had a large number of genes that were more than two-fold differentially expressed (Figure 3A, Supplementary table 3). Of note, DAF^lo^ GC B cells had upregulated genes, which suggested on-going somatic hypermutation, such as *BACH2*, *FOXO1* and *AICDA*^4,26–28^, whereas DAF^hi^ GC B cells showed elevated expression of genes involved in B cell differentiation (*PRDM1*, *IRF4*, *CCR6*), class switching (*BATF*), gene editing (*APOBEC3B*), and regulation of transcription (*MYC*) ^29–34^. Surprisingly, the gene encoding DAF (*CD55)*, showed similar transcription levels between the two subsets and we could confirm this by RT-qPCR (Figure 3B). Consistent with *PRDM1* being upregulated in the DAF^hi^ subset, we found that the gene product Blimp1 was similarly upregulated in the DAF^hi^ subset (Figure 3C). These data indicate that the CD19^+^CD20^+^IgD^−^CD38^+^DAF^hi^ B cell subset may comprise early GC-derived PB or PCs.

**Figure 3.**
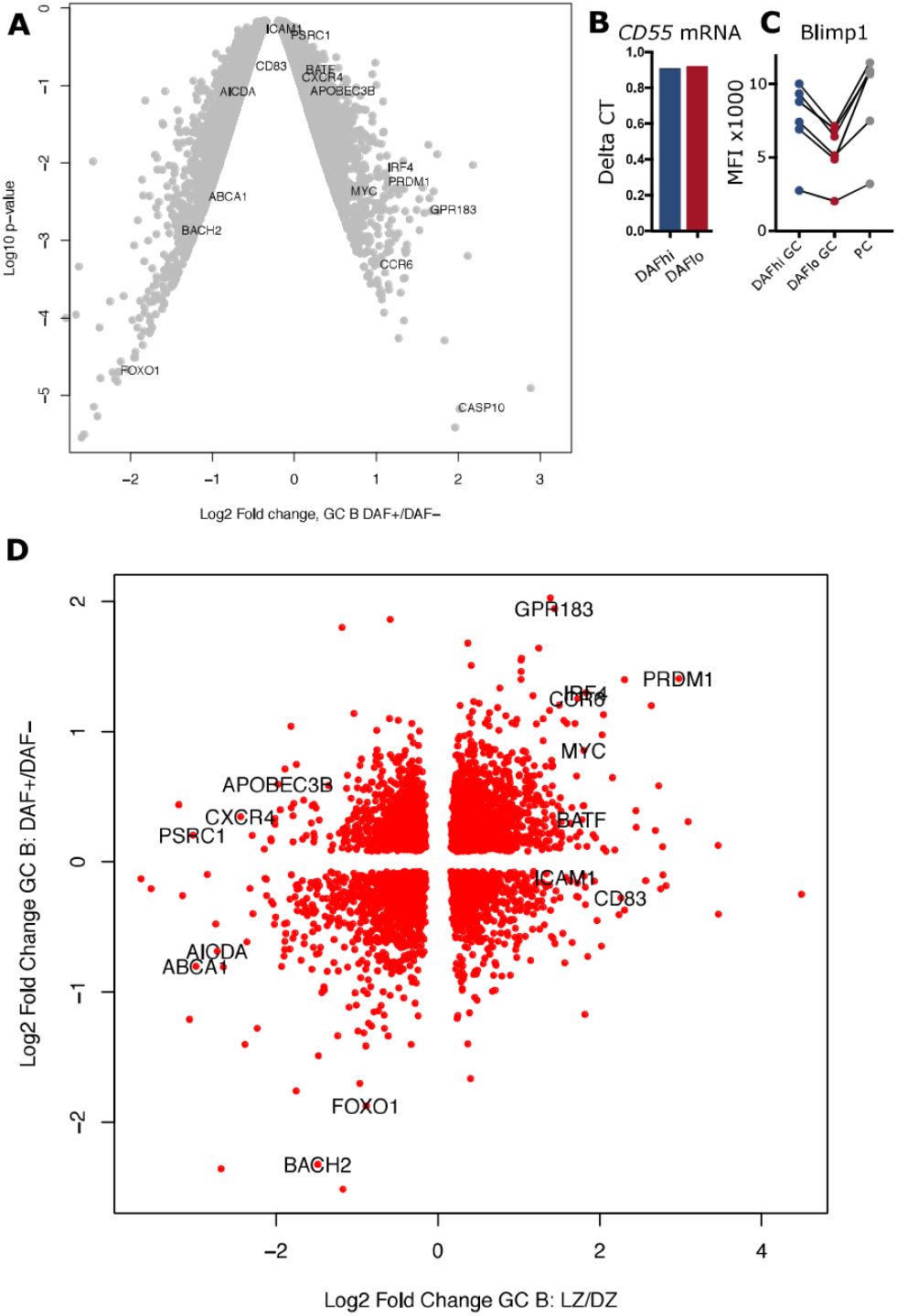
Transcriptomic analysis of DAF^hi^ and DAF^lo^ germinal center B cells. **(A)** Volcano plot showing differences in genes expressed by DAF^hi^ and DAF^lo^ GC B cells. **(B)** Delta CT of *CD55* gene expression, normalized to *ACTB*. **(C)** Flow cytometric analysis of Blimp1 expression on DAF^hi^, DAF^lo^ GC B cells and plasma cells (PC) (N = 6). **(D)** Clustering of DAF^hi^ and DAF^lo^ GC B cells in combination with transcriptomic data from sorted light zone (LZ) and dark zone (DZ) GC B cells. LZ and DZ data were obtained from Victora et al^35^.

Although many of the genes upregulated in DAF^lo^ GC B cells are typically associated with DZ B cells, the expression of the DZ marker CXCR4 was not increased. Instead, the LZ marker CD83 was increased in DAF^lo^ GC B cells and the DZ marker CXCR4 was increased in DAF^hi^ GC B cells. This indicated that both the LZ and DZ comprise DAF^hi^ and DAF^lo^ B cells. Therefore, we combined our transcriptional analysis of DAF^hi^ and DAF^lo^ GC B cells with published data on sorted LZ and DZ B cells^35^. Through this analysis, we found transcriptional patterns that applied to both DZ and LZ from sorted DAF^hi^ and DAF^lo^ GC B cells (Figure 3D). However, and consistent with our other data, the clustering suggested that DAF^lo^ B cells in the DZ are proliferating and undergoing SHM, whereas DAF^hi^ B cells contained a transcriptional profile indicative of final differentiation in the LZ.

### DAF expression in DZ and LZ of GCs

Next, we used expression of CXCR4 and CD83 to analyze DAF expression on GC B cells in DZ and LZ, respectively (Figure 4A). Consistent with our transcriptional analysis, we found DAF^hi^ and DAF^lo^ populations in both zones, where the frequency of DAF^hi^ cells was slightly decreased in the LZ (Figure 4B). It is established that B cells undergo apoptosis rather than lysis in GCs. We therefore assessed expression of the complement regulatory protein CD59 that inhibits formation of the membrane attack complex, and of complement receptors CD35 and CD21, which are involved in both complement regulation, and B cell activation. We found that CD59, CD35, and CD21 were expressed at similar levels regardless of zonal location or DAF phenotype, although we observed a trend of higher expression of CD35 and CD21 on DAF^hi^ cells in both zones. We could also confirm that GC B cells had increased expression of CD59, as compared to non-GC B cell subsets, excluding PBs and PCs (Figure 4D). In addition, the negative complement regulator CD46 (Figure 4E), which inactivates C4b and C3b, and the complement receptor 1, CD35 (Figure 3F), was also reduced in DAF^lo^ GC B cells. These data suggested that DAF^lo^ GC B cells may be primed for complement-dependent phagocytosis but not lysis.

**Figure 4.**
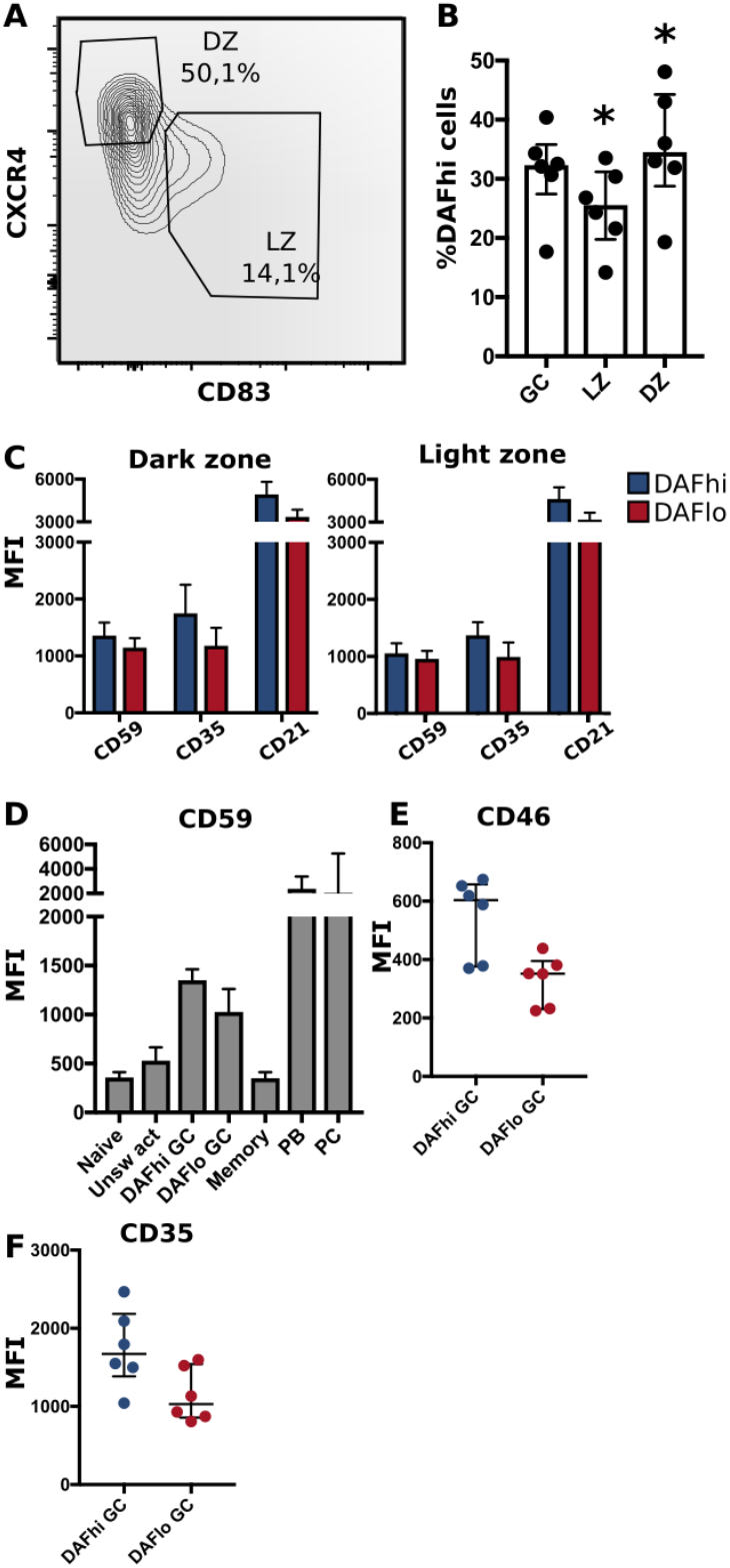
DAF expression on DZ and LZ B cells. **(A)** Representative gating strategy for light zone (LZ, CD83^+^ CXCR4^−^) and dark zone (DZ, CD83^−^ CXCR4^+^) of CD19^+^ CD20^+^ CD38^+^ IgD^−^ B cells. **(B)** Frequency of DAF^hi^ cells within the total GC B cell population and on LZ and DZ GC B cells. Bars represent median with interquartile range. *P < 0,05 by Wilcoxon signed-rank test. **(C)** Expression of CD59, CD35 and CD21 on DAF^hi^ or DAF^lo^ GC B cells in DZ (left) or LZ (right). **(D)** Expression of CD59 on naïve, unswitched activated (Unsw act), DAF^hi^ or DAF^lo^ GC B cells, memory B cells, plasmablasts (PB) and plasma cells (PC). **(E)** Expression of complement regulator CD46 on DAF^hi^ and DAF^lo^ GC B cells. **(F)** Expression of CD35 in DAF^hi^ and DAF^lo^ GC B cells. Median fluorescence intensity (MFI) is represented as individual dots where lines indicate median with interquartile range. Data was acquired from multiple patients (n = 6).

### B cell receptor stimulation leads to downregulation of DAF expression

Although downregulation of DAF was most pronounced on GC B cells, DAF was also reduced on a fraction of unswitched activated (CD38^+^IgD^−^) B cells in tonsils (Figure 2D). Hence, it was possible that BCR-stimulation alone or in concert with co-stimulatory factors could directly regulate DAF expression. To assess T cell-independent activation of B cells, we stimulated PBMCs from healthy donors with anti-IgM/IgG with or without CpG for four days (Figure 5A). We found that BCR-stimulation alone led to a reduction of DAF^hi^ B cells and that co-stimulation with CpG enhanced this reduction (Figure 5B). In contrast, CpG alone had no effect on the frequency of DAF^hi^ B cells. We proceeded to assess if T cell-dependent activation was similarly effective to enhance BCR-induced downregulation of DAF by co-incubating with anti-CD40, and recombinant IL-4 or IL-21, respectively (Figure 5C). Of these, IL-21 had a minor but significant effect on the frequency of DAF^hi^ B cells in the presence of anti-IgM/IgG (Figure 5D). These data demonstrated that DAF expression is regulated via BCR-stimulation, and that selected co-stimulatory factors during both T cell-dependent and T cell-independent activation of B cells enhanced the number of DAF^lo^ cells. Throughout all conditions used for T cell-dependent and independent stimulation of B cells, we consistently found that Blimp-1 was expressed at a higher level on DAF^hi^ than on DAF^lo^ B cells (Figure 5E). These data support our findings that DAF^hi^ GC B cells comprise a subpopulation of early GC-derived PB/PC. However, T cell-independent stimulation induced high level of Ki67 expression in both DAF^hi^ and DAF^lo^ B cells but T cell-dependent stimulation preferentially led to activation of DAF^lo^ B cells (Figure 5F). This demonstrated that the nature of co-stimulatory signals govern the activation state of DAF^hi^ or DAF^lo^ B cells, respectively.

**Figure 5.**
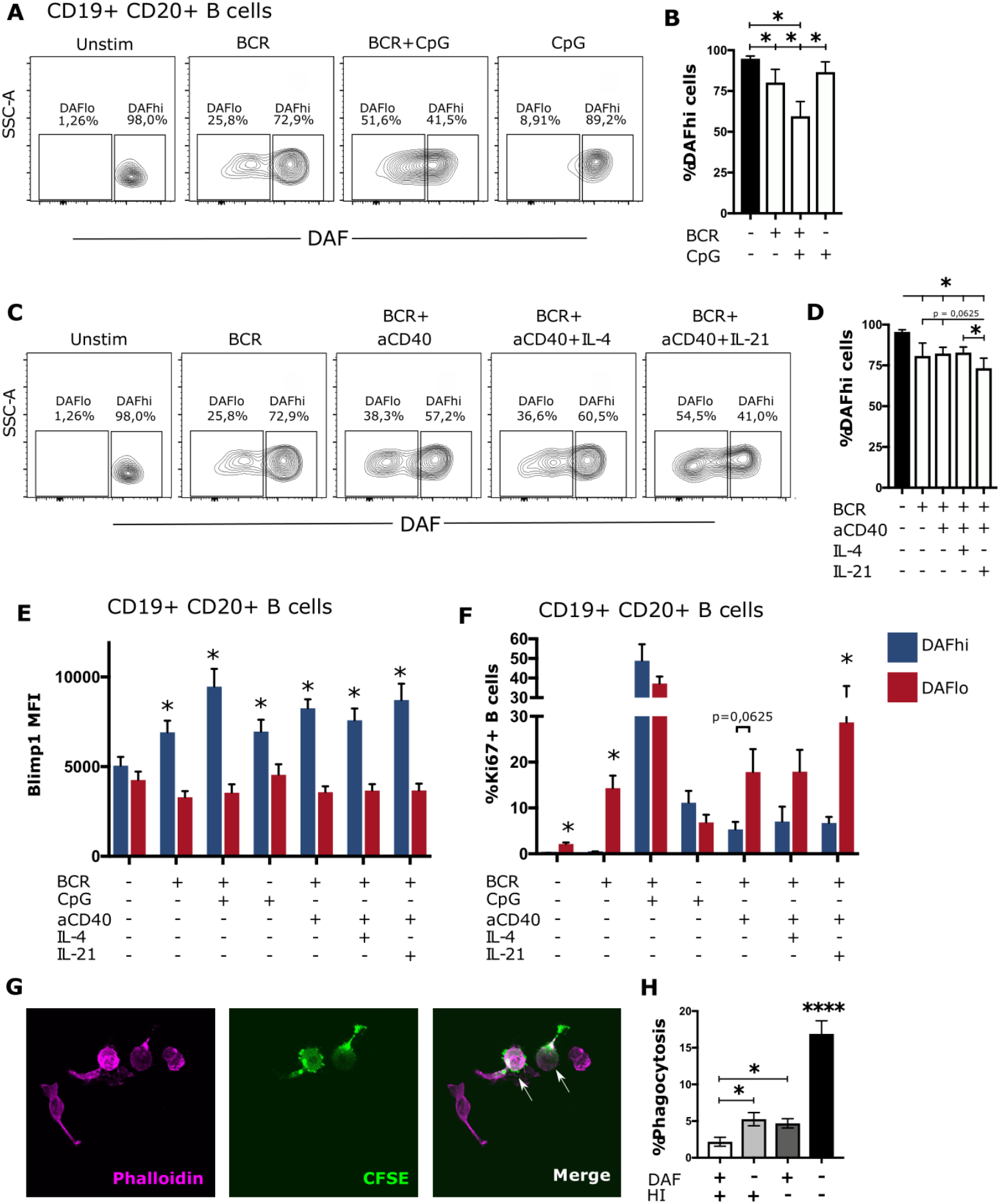
B cell receptor engagement leads to downregulation of DAF on B cells, in vitro. **(A)** Representative plots for discriminating DAF expression on CD19^+^ CD20^+^ B cells 4 days after stimulation with anti-IgM/IgG, CpG or a combination of both. **(B)** Quantification of data from (A). N=6. Median and interquartile range is shown. **(C)** Representative plots for discriminating DAF expression on CD19^+^ CD20^+^ B cells 4 days after stimulation with anti-IgM/IgG, anti-CD40, IL-4, IL-21 or a combination of these. **(D)** Quantification of data from (C). N=6. Median and interquartile range is shown. Intracellular expression of Blimp1 **(E)** or Ki67 **(F)** in DAF^hi^ and DAF^lo^ B cells after 4 day stimulation with anti-IgM/IgG, CpG, anti-CD40, IL-4, IL-21 or a combination of these. N=6. Median and interquartile range is shown. *P < 0,05 by Wilcoxon signed-rank test. **(G)** Representative image of phalloidin labelled primary human macrophages (magenta), CFSE stained sorted B cells (green) and phagocytosed B cells (magenta and green). **(F)** Quantification of phagocytosed B cells after co-incubation of sorted and CFSE labelled DAF^hi^ and DAF^lo^ GC B cells with human serum before or after heat inactivation (HI). 200 macrophages were counted per slide and cells double positive for CFSE and phalloidin were considered phagocytosing. Data from 3 independent experiments is shown (GC B cells from three unique individuals). Mean and standard error of mean is shown. *P < 0,05; **P < 0,01; ****P < 0,0001 by Mann-Whitney test.

### Phagocytosis of DAF^lo^ GC B cells is enhanced in the presence of complement

A previous study demonstrated that DAF-deficient human T cells accumulate C3d on their cell surfaces *in vitro*^20^, and that this can facilitate phagocytosis^36^. Therefore, we hypothesized that DAF^lo^ GC B cells would be more efficiently phagocytosed than their DAF^hi^ counterparts, in a complement-dependent manner. To test this hypothesis, we sorted CFSE-labeled DAF^hi^ and DAF^lo^ GC B cells and co-cultivated these with primary human macrophages in the presence of human serum prior or after heat inactivation of complement. Phagocytosis of B cells was then assessed by counting Phalloidin^+^ macrophages that had internalized vesicles that contained the CFSE dye from the sorted B cells. (Figure 5G). Consistent with our hypothesis, we could show efficient and complement-dependent phagocytosis of DAF^lo^ GC B cells, in comparison with DAF^hi^ GC B cells (Figure 5H). Together, these observations strongly suggest that GC B cells are pre-primed for complement-dependent phagocytosis, and that this is regulated by a reduction of DAF on their cell surface.

### Modulation of DAF expression during B cell hematopoiesis in the bone marrow

Similar to GC structures in secondary lymphoid organs, the bone marrow represents another site of extensive B cell proliferation. To understand if DAF may play a role also in early B cell development, we obtained human bone marrow samples from routine biopsies and stratified the stages of B cell development by flow cytometry^37^. Briefly, we separated PCs (CD19^+^ CD38^hi^) from other B cells and precursors (CD19^+^ CD38^dim/lo^) (Figure 6A). Pro-B cells were identified as CD34^+^ CD10^+^ (Figure 6B). Then, we used CD20 and CD10 identify Pre-B1 cells (CD10^+^ CD20^−^), pre-B2 cells (CD10^+^ CD20^+^), transitional B cells (CD10^dim^ CD20^+^)and mature B cells (CD10^−^ CD20^+^) (Figure 6C). The pre-B2 subset could then be further divided into large and small cells, based on forward side scatter. Subsequent assessment of DAF on the different B cell populations revealed that surface expression of DAF was low from the Pro-B stage until the large immature stage, where the expression followed a bimodal pattern (Figure 6D). Small pre-B2 cells showed uniformly low expression of DAF whereas large pre-B2 cells showed a bimodal pattern of DAF expression. From the transitional B cell subset and onward, DAF expression was uniformly high, where PCs demonstrated the highest surface expression of DAF. Both MFI and %DAF^hi^ cells of parent followed this pattern throughout the developmental stages (Figure 6E, 6F). Together, these data demonstrate that DAF is upregulated on a fraction of cells at a late developmental stage where testing of the pre-BCR or formation of a functional BCR occurs. These data demonstrate that, similar to GCs, DAF is regulated also during B cell development in the bone marrow.

**Figure 6.**
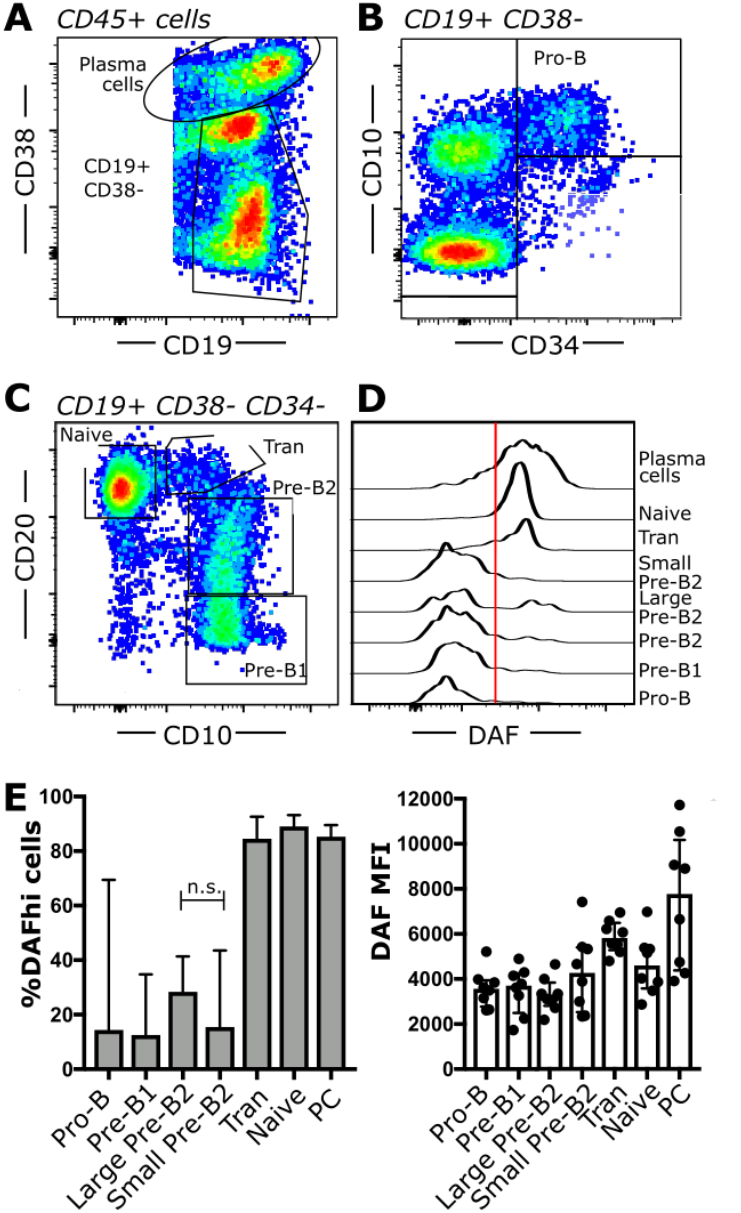
Regulation of DAF expression during B cell hematopoiesis. **(A-C)** Flow cytometric analysis of human bone marrow aspirates. Shown are **(A)**PCs (CD19^+^ CD38^hi^) and B cells (CD19^+^ CD38^dim/lo^), **(B)**Pro-B cells (CD34^+^ CD10^+^) and **(C)**Pre-B1 cells (CD10^+^ CD20^−^), pre-B2 cells (CD10^+^ CD20^+^), transitional B cells (CD10^dim^ CD20^+^) and mature B cells (CD10^−^ CD20^+^). **(D)** Representative histogram of DAF expression from respective B cell subset in bone marrow. The line indicates the cut-off for determination of high or low DAF expression. **(E)** Quantification of data from (D). Shown is median with interquartile range. n.s. non-significant; *P < 0,05; **P < 0,01. N = 8.

## Discussion

The complement regulatory protein DAF is well known to inhibit complement activation on cell surfaces. Here, we demonstrate that human GC B cells downregulate DAF on their cell surfaces, and that one function of this downregulation is to prime GC B cells for complement-dependent phagocytosis.

Our transcriptional data suggest that the distribution of DAF^lo^ and DAF^hi^ GC B cells are located in both the DZ and LZ of the GC structure. Based on the preferential transcription of *AICDA*, *FOXO1* and *BACH2*, DAF^lo^ GC B cells in the DZ undergo somatic hypermutation. Moreover, we could also decipher that DAF^hi^ GC B cells that are located in the LZ comprise cells that express *PRDM1* and *IRF4*, that are associated with PC differentiation. This was corroborated by upregulation of Blimp1 also on the protein level. These data suggest that DAF may be a useful marker to further study early PC differentiation in GCs.

Our *in vitro* experiments demonstrated that B cells downregulate surface expression of DAF after BCR engagement alone. This is consistent with our data that show downregulation of DAF on fractions of unswitched activated or memory B cells in circulation and in tonsils. We initially had a hypothesis that DAF would be upregulated on GC B cells that undergo successful selection, but addition of anti-CD40 to mimic interaction with T cells did only minorly affect DAF expression, and then only in concert with addition of IL-21. Instead, we found that addition of anti-CD40, IL4 or IL21 preferentially led to an upregulation of Ki67 on DAF^lo^ B cells. Since entry of activated B cells into GC occurs via a T cell dependent checkpoint^38^, this could explain why 80-90% of all GC B cells are DAF^lo^. However, during the on-going GC reaction, DAF expression may be regulated largely independent from T cell-dependent selection. In contrast, CpG lead to upregulation of Ki67 on both DAF^hi^ and DAF^lo^ B cells. It is therefore possible that T cell-dependent or independent responses have different requirements for modulation of DAF after activation. In this study, we did not find an overall downregulation of the *CD55* gene on cells with low surface expression of DAF. This suggests that surface expression of DAF is regulated either by post-translational cleavage or by alternative splicing^39^ and studies are on-going to clarify the regulation of DAF.

The complement system is involved in both innate and adaptive immune responses but it also facilitates the removal of dead or dying cells via a non-inflammatory process^40^. Products of cleaved complement protein C3, such as C3b and C3d, are involved in this process^18^. The abundance of C3d in GCs is partially explained by the critical role of C3d for transport of immune complexes into lymphoid follicles and activation of antigen-specific follicular B cells^41^. However, data from several early investigations has suggested that GCs may be subject to a local complement cascade, including attachment of complement to cells ^42,43^. The low DAF expression on GC B cells may explain this, as this would allow for attachment of C3b on cell surfaces^18^. Since our flow cytometric data was generated after gating on viable cells (Supplementary Figure S2), they demonstrate that downregulation of DAF had not led to apoptosis, nor lysis via the membrane-attack complex. This latter may be explained by consistent expression of CD59 on GC B cells to hinder the generation of the membrane attack complex by inhibition of C5 convertase^43^. It has been described that regulation of complement receptors on GC B cells can influence the threshold for BCR-mediated activation^24^, enhance antigen uptake^45^, and also facilitate the attachment of these fragments onto B cells^46^. We also found that DAF expression was modulated during B cell hematopoiesis in the bone marrow. This opens up a possibility that regulation of DAF may serve a similar function during B cell development as we show for GC B cells; to prime cells for phagocytosis.

Here, we demonstrate a role of DAF for phagocytosis of GC B cells. While we did not investigate if modulation of DAF could have other beneficial functions for GC B cells, DAF deficient mouse macrophages and dendritic cells have been shown to present antigen more efficiently to T cells than their DAF expressing counterparts^47^. It is therefore possible that the regulation of DAF may serve a dual role where it also allows activated GC B cells to more easily interact with T cells. This would also be in line with previous observations that complement interaction facilitates antigen uptake in B cells^45^.

Collectively, our data demonstrates a novel role of DAF for regulation of phagocytosis of GC B cells and that modulation of DAF may also play an important role during B cell development. This may explain how B cell homeostasis is maintained at locations where extensive proliferation and apoptosis occurs.

## Materials and methods

### Donors and tissues

The research was carried out according to The Code of Ethics of the World Medical Association (Declaration of Helsinki). Ethical permits were obtained from the Swedish Ethical review authority (No: 2016/53-31, 04-113M, 07-162M and 2014/233) and all samples were collected after receiving informed consent from patient or patient’s guardian. Briefly, blood was collected in EDTA tubes and PBMCs were isolated using a Ficoll-Paque density gradient centrifugation. Tonsillar cell suspensions were prepared by tissue homogenizing in RPMI-1640 medium and passed through a 70 μm cell strainer. Red blood cells were lysed using BD PharmLyse lysis buffer according to manufacturer’s instructions. PBMCs from healthy donors were isolated by Ficoll-Paque density gradient from buffy coats from routine blood donations at the Blood Central at Umeå University Hospital, Umeå, Sweden. All cell suspensions except bone marrow aspirates were frozen in fetal bovine serum (FBS) (Gibco) with 10% DMSO and stored in liquid N_2_. Bone marrow aspirates were obtained from routine sampling at the Department of Pathology, Umeå University Hospital.

### Flow cytometry

Antibodies used are listed in supplementary table 1. Frozen suspensions of PBMCs and tonsils were thawed, washed and resuspended in PBS with 2% FBS, then stained with Fixable Viability Stain 780 (BD Biosciences), followed by antibody staining for 30 minutes at 4°C. Intracellular staining for transcription factors was performed using the eBioscience FoxP3/Transcription Factor Staining Buffer set according to manufacturer’s instruction (ThermoFisher). Cells were acquired on a BD LSRII or BD FACSAria III. Cell sorting was done on BD FACSAria III. Bone marrow samples were processed by routine diagnostic procedures and acquired on a BD FACSCanto II. All data were analyzed using the FlowJo v10 software.

### Tissue immunofluorescence

Tonsils were fixed for 4 hours in PBS + 4% paraformaldehyde, then incubated overnight in 30% sucrose. Samples were embedded in OCT (HistoLab) and stored at −80°C. 20 μm sections of the tissues were cut in a cryostat. The sections were blocked for 1 hour at room temperature in PBS + 5% FBS + 0.1% Triton, then stained with antibodies against CD19, IgD, CXCR4 and DAF. Full details of antibodies are listed in supplementary table 2. Stained sections were imaged on a Zeiss LSM 710 confocal microscope with 405, 488, 561 and 647 nm laser lines, using a Plan Apochromat 20x objective.

### Cell culture

PBMCs from healthy donors were seeded at 1×10^6^ cells/ml in a 96-well plate containing RPMI-1640, L-Glutamine (Gibco), 10% fetal bovine serum (Gibco) and 100 U/L Penicillin-Streptomycin (Gibco). Cells were then stimulated with 10 μg/ml goat-anti human IgM+IgG (Jackson Laboratories), 2.5 μM CpG B (ODN 2006, Invivogen), 1 μg/ml anti-CD40 (G28.5, Abcam), 25 ng/ml IL-4 (Abcam) or 25 ng/ml IL-21 (Abcam). All incubations were at 37°C, 5%CO2.

### Microarray

DAF^hi^ and DAF^lo^ GC B cells (CD19^+^ CD20^+^ CD38^+^ IgD^−^) were resuspended in RLT cell lysis buffer (Qiagen) after flow cytometric sorting. Total RNA was extracted (Qiagen) and microarray was performed using Affymetrix Human Clariom D microarrays (Bioinformatics and Expression Analysis core facility at Karolinska Institutet, Huddinge, Sweden). Data were analyzed in R. First the data was RMA normalized. Next, limma was used to solve the differential expression regression problems using empirical Bayes. In all cases we regressed out donor effects (~x+donor, where x is e.g. DAF^hi^ vs DAF^lo^).

### RT-qPCR of DAF expression

5000 each of DAF^hi^ and DAF^lo^ GC B cells were resuspended in RLT buffer after FACS sorting. RNA was extracted using Qiagen RNEasy Micro Kit according to instructions. One-step RT-qPCR was perfomed with LightCycler 480 RNA Master Hydrolysis Probes (Roche) and commercially available *CD55* and *ACTB* TaqMan primers and probes during 5 min at 60°C, 1 min at 95°C, then 15s 95°C and 1 min 60°C for 45 cycles. RNA was loaded in triplicates and the reaction was run on a QuantStudio 5 Real-Time PCR System machine. *CD55* expression was normalized to ACTB expression to obtain the delta Ct values.

### Phagocytosis assay

Primary human macrophages were cultivated from purified PBMCs from a healthy donor. 1.5 million PBMCs/well were plated in RPMI-1640 supplemented with 10% FBS and 1% Pen-Strep on 13 mm circular coverslips in a 24-well plate for 2h. Non-adherent cells were rinsed off with PBS, and RPMI-1640 supplemented as described and with additional 20 mM Hepes and 25 ng/ml M-CSF (R&D Systems). Medium was changed every third day and the cells were allowed to differentiate for 10 days. Normal human serum was collected and pooled from 6 healthy donors and stored at −80°C immediately after isolation. After a 1h incubation of sorted DAF^lo^ or DAF^hi^ GC B cells with macrophages in RPMI-1640, supplemented with 10μM CaCl_2_ and 10μM MgCl_2_ and 10% of either thawed or heat-inactivated human serum, samples were fixed with 4% paraformaldehyde in PBS and permeabilized with 0.1% Triton X-100, followed by staining with AlexaFluor-546 phalloidin (ThermoFisher). Phagocytosis was quantified by microscopy where phalloidin^+^ macrophages were counted, and macrophages containing CFSE signal and phagocytic vesicles were considered as phagocytosing. A total of 200 macrophages per well were counted.

### Statistics

All statistic calculations were performed using GraphPad Prism 7. For comparisons between populations within the same patient, we performed Wilcoxon matched-pairs signed rank test. For the comparisons between different groups, we used the Mann-Whitney test. P-values lower than 0.05 were considered as significant.

## Data availability

The microarray data is available under accession GEO (GSE153741). The R code is available at Github (**https://github.com/henriksson-lab/mattias-daf**).

## Authorship

### Contributions

A.D. designed and performed experiments and wrote the manuscript. J.L. assisted with experimental work and critically read the manuscript. A.H. organized tonsillectomies and provided expertise in sample processing, and critically read the manuscript. J.M. provided conceptual suggestions, assisted with microarray, and critically read the manuscript. C.A. provided HFRS samples and supervised the project. J.H. assisted with bioinformatic analysis and critically read the manuscript. M.H. arranged bone marrow sampling and analysis, provided expertise in B cell development, and critically read the manuscript. M.N.E.F. conceived and supervised the study, designed experiments, and wrote the manuscript.

## Acknowledgements

The authors thank Remigius Gröning for commenting on the manuscript; Ellen Ernhill, MD and Anders Erlandsson, MD, for providing tonsils; Elin Arvidsson for processing and acquisition of the bone marrow samples; Myriam Martin and Anna Blom at Lunds University for constructive discussions with regards to complement activation. We also would like to thank the Affymetrix core facility at Novum, BEA, Bioinformatics and Expression Analysis, which is supported by the board of research at the Karolinska Institute and the research committee at the Karolinska hospital.

## References

1. Hozumi N, Tonegawa S. Evidence for somatic rearrangement of immunoglobulin genes coding for variable and constant regions. 1976 [classical article]. J Immunol. 2004;173(7):4260–4264.

2. Talmage DW. Allergy and immunology. Annu Rev Med. 1957;8:239–256.

3. Burnet FM. A modification of Jerne's theory of antibody production using the concept of clonal selection. CA Cancer J Clin. 1976;26(2):119–121.

4. Muramatsu M, Kinoshita K, Fagarasan S, Yamada S, Shinkai Y, Honjo T. Class switch recombination and hypermutation require activation-induced cytidine deaminase (AID), a potential RNA editing enzyme. Cell. 2000;102(5):553–563.

5. McKean D, Huppi K, Bell M, Staudt L, Gerhard W, Weigert M. Generation of antibody diversity in the immune response of BALB/c mice to influenza virus hemagglutinin. Proc Natl Acad Sci U S A. 1984;81(10):3180–3184.

6. Berek C, Berger A, Apel M. Maturation of the immune response in germinal centers. Cell. 1991;67(6):1121–1129.

7. Neuberger MS, Lanoue A, Ehrenstein MR, Batista FD, Sale JE, Williams GT. Antibody diversification and selection in the mature B-cell compartment. Cold Spring Harb Symp Quant Biol. 1999;64:211–216.

8. Victora GD, Nussenzweig MC. Germinal centers. Annu Rev Immunol. 2012;30:429–457.

9. Green DR, Oguin TH, Martinez J. The clearance of dying cells: table for two. Cell Death Differ. 2016;23(6):915–926.

10. Arandjelovic S, Ravichandran KS. Phagocytosis of apoptotic cells in homeostasis. Nat Immunol. 2015;16(9):907–917.

11. Domen J, Cheshier SH, Weissman IL. The role of apoptosis in the regulation of hematopoietic stem cells: Overexpression of Bcl-2 increases both their number and repopulation potential. J Exp Med. 2000;191(2):253–264.

12. Mayer CT, Gazumyan A, Kara EE, et al. The microanatomic segregation of selection by apoptosis in the germinal center. Science. 2017;358(6360).

13. Flemming W. Studien über Regeneration der Gewebe. Arch Mikr Anat. 1885;24:50.

14. Swartzendruber DC, Congdon CC. Electron microscope observations on tingible body macrophages in mouse spleen. J Cell Biol. 1963;19(3):641–646.

15. Smith JP, Burton GF, Tew JG, Szakal AK. Tingible body macrophages in regulation of germinal center reactions. Dev Immunol. 1998;6(3-4):285–294.

16. Sato K, Honda SI, Shibuya A, Shibuya K. Cutting Edge: Identification of Marginal Reticular Cells as Phagocytes of Apoptotic B Cells in Germinal Centers. J Immunol. 2018;200(11):3691–3696.

17. Liu YJ, Joshua DE, Williams GT, Smith CA, Gordon J, MacLennan IC. Mechanism of antigen-driven selection in germinal centres. Nature. 1989;342(6252):929–931.

18. Martin M, Blom AM. Complement in removal of the dead - balancing inflammation. Immunol Rev. 2016;274(1):218–232.

19. Eldewi DM, Alhabibi AM, El Sayed HME, et al. Expression levels of complement regulatory proteins (CD35, CD55 and CD59) on peripheral blood cells of patients with chronic kidney disease. Int J Gen Med. 2019;12:343–351.

20. Ozen A, Comrie WA, Ardy RC, et al. CD55 Deficiency, Early-Onset Protein-Losing Enteropathy, and Thrombosis. N Engl J Med. 2017;377(1):52–61.

21. Takeda J, Miyata T, Kawagoe K, et al. Deficiency of the GPI anchor caused by a somatic mutation of the PIG-A gene in paroxysmal nocturnal hemoglobinuria. Cell. 1993;73(4):703–711.

22. Richards SJ, Morgan GJ, Hillmen P. Immunophenotypic analysis of B cells in PNH: insights into the generation of circulating naive and memory B cells. Blood. 2000;96(10):3522–3528.

23. Jonsson CB, Figueiredo LT, Vapalahti O. A global perspective on hantavirus ecology, epidemiology, and disease. Clin Microbiol Rev. 2010;23(2):412–441.

24. Cherukuri A, Cheng PC, Sohn HW, Pierce SK. The CD19/CD21 complex functions to prolong B cell antigen receptor signaling from lipid rafts. Immunity. 2001;14(2):169–179.

25. Kischkel FC, Hellbardt S, Behrmann I, et al. Cytotoxicity-dependent APO-1 (Fas/CD95)-associated proteins form a death-inducing signaling complex (DISC) with the receptor. Embo j. 1995;14(22):5579–5588.

26. Miura Y, Morooka M, Sax N, et al. Bach2 Promotes B Cell Receptor-Induced Proliferation of B Lymphocytes and Represses Cyclin-Dependent Kinase Inhibitors. J Immunol. 2018;200(8):2882–2893.

27. Muto A, Tashiro S, Nakajima O, et al. The transcriptional programme of antibody class switching involves the repressor Bach2. Nature. 2004;429(6991):566–571.

28. Inoue T, Shinnakasu R, Ise W, Kawai C, Egawa T, Kurosaki T. The transcription factor Foxo1 controls germinal center B cell proliferation in response to T cell help. J Exp Med. 2017;214(4):1181–1198.

29. Turner CA, Jr., Mack DH, Davis MM. Blimp-1, a novel zinc finger-containing protein that can drive the maturation of B lymphocytes into immunoglobulin-secreting cells. Cell. 1994;77(2):297–306.

30. Sciammas R, Davis MM. Modular nature of Blimp-1 in the regulation of gene expression during B cell maturation. J Immunol. 2004;172(9):5427–5440.

31. Calado DP, Sasaki Y, Godinho SA, et al. The cell-cycle regulator c-Myc is essential for the formation and maintenance of germinal centers. Nat Immunol. 2012;13(11):1092–1100.

32. Ise W, Kohyama M, Schraml BU, et al. The transcription factor BATF controls the global regulators of class-switch recombination in both B cells and T cells. Nat Immunol. 2011;12(6):536–543.

33. Suan D, Kräutler NJ, Maag JLV, et al. CCR6 Defines Memory B Cell Precursors in Mouse and Human Germinal Centers, Revealing Light-Zone Location and Predominant Low Antigen Affinity. Immunity. 2017;47(6):1142–1153.e1144.

34. Zou J, Wang C, Ma X, Wang E, Peng G. APOBEC3B, a molecular driver of mutagenesis in human cancers. Cell Biosci. 2017;7:29.

35. Victora GD, Dominguez-Sola D, Holmes AB, Deroubaix S, Dalla-Favera R, Nussenzweig MC. Identification of human germinal center light and dark zone cells and their relationship to human B-cell lymphomas. Blood. 2012;120(11):2240–2248.

36. Gaither TA, Vargas I, Inada S, Frank MM. The complement fragment C3d facilitates phagocytosis by monocytes. Immunology. 1987;62(3):405–411.

37. Theunissen P, Mejstrikova E, Sedek L, et al. Standardized flow cytometry for highly sensitive MRD measurements in B-cell acute lymphoblastic leukemia. Blood. 2017;129(3):347–357.

38. Schwickert TA, Victora GD, Fooksman DR, et al. A dynamic T cell-limited checkpoint regulates affinity-dependent B cell entry into the germinal center. J Exp Med. 2011;208(6):1243–1252.

39. Vainer ED, Meir K, Furman M, Semenenko I, Konikoff F, Vainer GW. Characterization of novel CD55 isoforms expression in normal and neoplastic tissues. Tissue Antigens. 2013;82(1):26–34.

40. Kerr JF, Wyllie AH, Currie AR. Apoptosis: a basic biological phenomenon with wide-ranging implications in tissue kinetics. Br J Cancer. 1972;26(4):239–257.

41. Carroll MC, Isenman DE. Regulation of humoral immunity by complement. Immunity. 2012;37(2):199–207.

42. Feucht HE, Jung CM, Gokel MJ, et al. Detection of both isotypes of complement C4, C4A and C4B, in normal human glomeruli. Kidney Int. 1986;30(6):932–936.

43. Gajl-Peczalska KJ, Fish AJ, Meuwissen HJ, Frommel D, Good RA. Localization of immunological complexes fixing beta1C (C3) in germinal centers of lymph nodes. J Exp Med. 1969;130(6):1367–1393.

44. Rollins SA, Zhao J, Ninomiya H, Sims PJ. Inhibition of homologous complement by CD59 is mediated by a species-selective recognition conferred through binding to C8 within C5b-8 or C9 within C5b-9. J Immunol. 1991;146(7):2345–2351.

45. Brimnes MK, Hansen BE, Nielsen LK, Dziegiel MH, Nielsen CH. Uptake and presentation of myelin basic protein by normal human B cells. PLoS One. 2014;9(11):e113388.

46. Nielsen CH, Pedersen ML, Marquart HV, Prodinger WM, Leslie RG. The role of complement receptors type 1 (CR1, CD35) and 2 (CR2, CD21) in promoting C3 fragment deposition and membrane attack complex formation on normal peripheral human B cells. Eur J Immunol. 2002;32(5):1359–1367.

47. Fang C, Miwa T, Song WC. Decay-accelerating factor regulates T-cell immunity in the context of inflammation by influencing costimulatory molecule expression on antigen-presenting cells. Blood. 2011;118(4):1008–1014.

