## Supplemental figures for "Regulation of Decay Accelerating Factor primes human germinal center B cells for phagocytosis"

### Figure S1

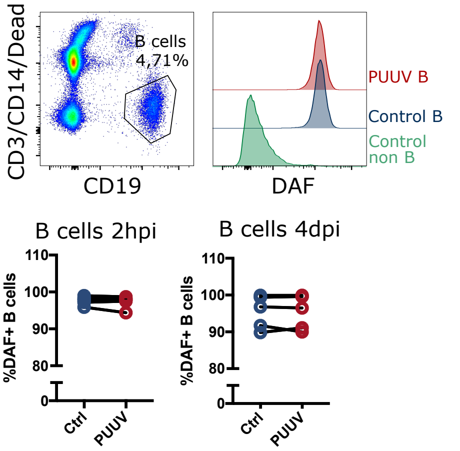


#### Supplemental figure S1.

In vitro incubation of PBMCs with PUUV Hantavirus and expression of DAF post-incubation

### Figure S2

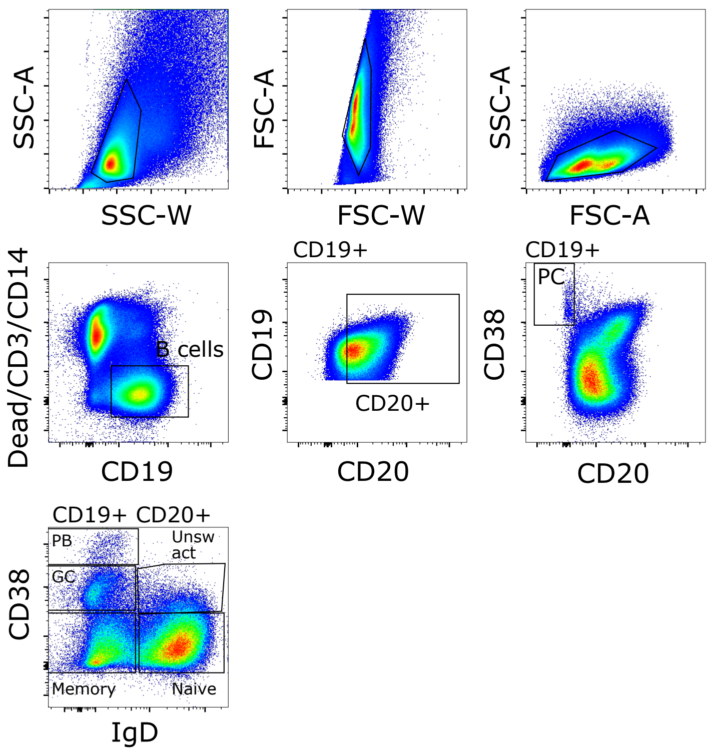


Supplementary figure S2.

Flow cytometric gating of tonsils and PBMCs.
